# The invasion of de-differentiating cancer cells into hierarchical tissues

**DOI:** 10.1101/574251

**Authors:** Da Zhou, Yue Luo, David Dingli, Arne Traulsen

**Affiliations:** School of Mathematical Sciences and Fujian Provincial Key Laboratory of Mathematical Modeling and High-Performance Scientific Computation, Xiamen University, Xiamen 361005, People’s Republic of China; Department of Evolutionary Theory, Max Planck Institute for Evolutionary Biology, August-Thienemann-Str. 2, 24306 Plön, Germany; Division of Hematology and Department of Internal Medicine, Mayo Clinic, 2nd St SW, 55905 Rochester, MN, USA

**Author notes:** (DZ); (AT).

## Abstract

Many fast renewing tissues are characterized by a hierarchical cellular architecture, with tissue specific stem cells at the root of the cellular hierarchy and differentiating into a whole range of specialized cells. There is increasing evidence that tumors are structured in a very similar way, mirroring the hierarchical structure of the host tissue. In some tissues, differentiated cells can also revert to the stem cell phenotype, which increases the risk that cells that have already acquired mutations lead to long lasting clones in the tissue. Recently, the modelling community has paid special attention to the consequences of de-differentiation on cellular hierarchies. However, the adaptive significance of de-differentiation is still poorly understood and thus it is unclear under which circumstances de-differentiating cells will invade a tissue. To address this, we developed mathematical models to investigate how de-differentiation could be selected as an adaptive mechanism in the context of cellular hierarchies. We consider the cases of stepwise and jumpwise de-differentiation in this study. Our results show that the emergence of de-differentiation is driven by the combination of the properties of the cellular hierarchy and the de-differentiation pattern and derive thresholds for which de-differentiation is expected to emerge.

## Introduction

In multicellular organisms, it is important that the inevitable replication errors of cells do not persist and threaten the functioning of the organism as a whole. Many tissues that need to undergo continuous cell turnover are organized in a hierarchical multi-compartment structure, which reduces the risk of the persistence of such mutations [1–13]. Each compartment represents a certain stage of cellular differentiation (Fig 1). At the root of the cellular hierarchy are tissue specific stem cells (SCs) which are capable of self-renewal and differentiation into more mature cells [14]. It is often argued that cancers may have similar hierarchical structures, where cancer stem cells (CSCs) possess characteristics associated with SCs in normal tissues [14, 15]. The CSCs scenario assumes that some cancerous tissues are hierarchically organized, similar to normal tissues [16].

**Figure 1:**
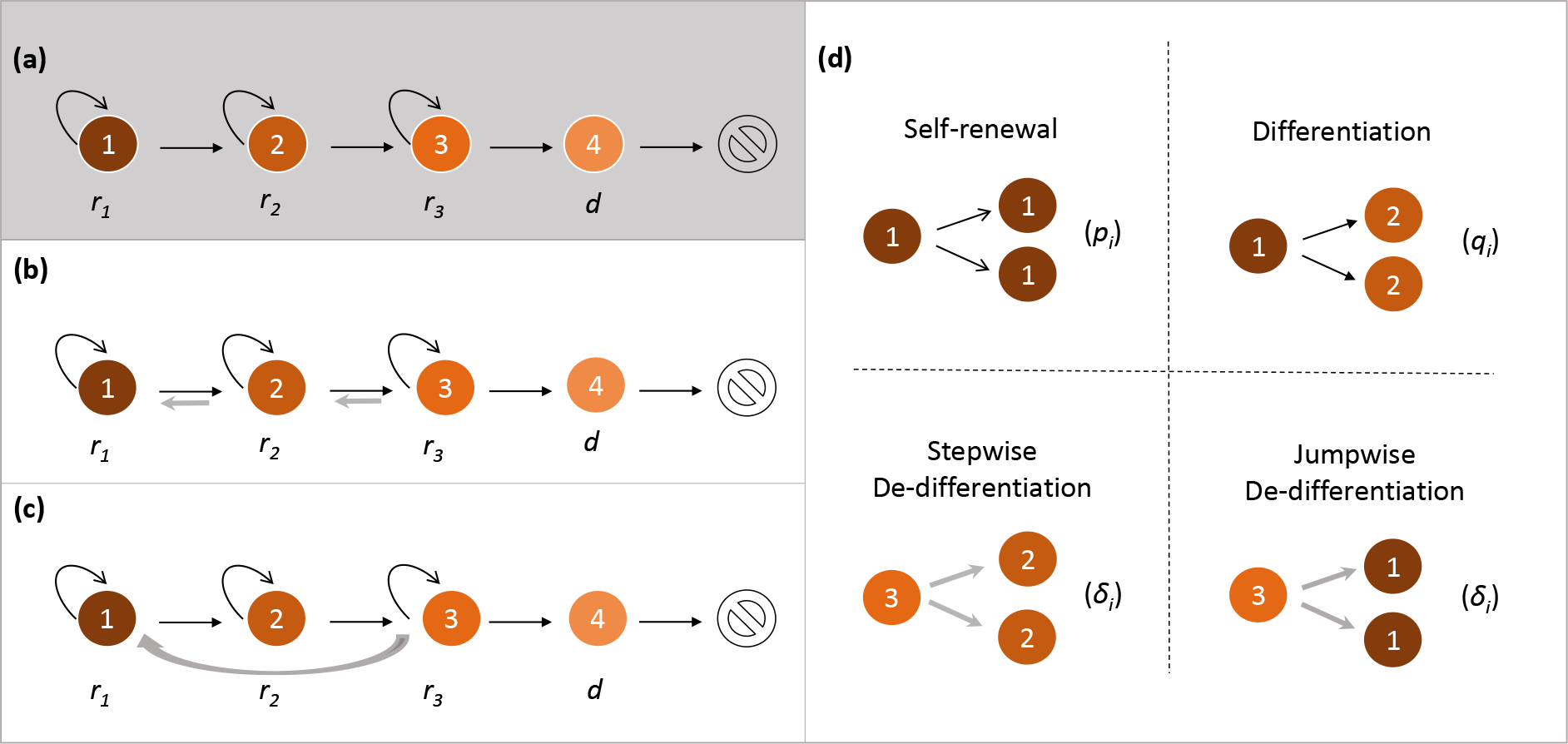
Representation of our models. We illustrate our models by considering a four-compartment hierarchical structure. **(a)** Null model without de-differentiation. Each compartment represents a certain stage of cell differentiation. For example, compartment 1 represents stem cell which performs cell division with rate *r*_1_. In each cell division, it can either give birth to two identical stem cells (self-renewal) or two identical daughter cells in adjacent downstream compartment 2 (differentiation). Similar division pattern can also happen to cells in compartments 2 and 3 (with division rates *r*_2_ and *r*_3_ respectively). Compartment 4 represents terminally differentiated cells which cannot divide and are removed from the tissue at rate d. **(b)** Stepwise de-differentiation case. Based on the hierarchical structure, we consider de-differentiation from downstream compartment *i* + 1 to adjacent upstream compartment *i*. (c) Jumpwise de-differentiation case, in which de-differentiation happens directly from compartment 3 to 1 without cells reaching the state in compartment 2. **(d)** The four cell division patterns used in our models.

The hierarchical tissue architecture proposes a unidirectional cascade from less differentiated stages to more differentiated stages (Fig 1 a). This would minimize the risk of the accumulation of genomic damage in the long term self renewing stem cells. However, there is significant evidence that the directional relation between different stages of differentiation could be broken in some tissues [17–22]. In other words, cells in later differentiated stages can, under some circumstances, revert to earlier differentiated stages, or even the stem cell stage, in a process known as de-differentiation (Fig 1b and c). De-differentiation could play an important role in regeneration and tumorigenesis [17]. In particular, even though the origin of cancer stem cells is still an open question, growing evidence shows that non-stem cancer cells can reacquire stem-like characteristics in colon cancer [23], breast cancer [20, 21], melanoma [24], leukemia [25–28], glioblastoma [29], and other cancers. For example, expression of the MLL-AF9 gene in committed hematopoietic progenitor cells led to the development of a leukemic stem cell population where only four of these cells were able to result in disease in a mouse model that could be transferred from one mouse to another, confirming the presence of a stem cell population [27].

More recently, special attention has been paid to the effect of de-differentiation on the cellular hierarchy by mathematical modeling of their impact. Previous works have considered how de-differentiation influences the waiting time to carcinogenesis [30], the fixation probability of a mutant [31, 32], the phenotypic equilibrium [33–35], transient overshoots [36, 37], and radiation sensitivity [29]. However, the adaptive significance of de-differentiation is still poorly understood: Under which circumstances would de-differentiation arise in the first place and rise in abundance? It is still unclear whether de-differentiation is a crucial improvement or just an unintended consequence of cellular hierarchy. A related problem is why de-differentiation arises in only some tumors, but not in others. Moreover, the comparison between different patterns of de-differentiation has received little attention.

Here we develop a matrix population model [38] of a stage-structured population for studying the evolution of de-differentiation. Two typical de-differentiation cases are taken into account in our model: One is stepwise de-differentiation which happens from a downstream compartment to an adjacent upstream compartment (Fig 1b), the other is jumpwise de-differentiation which is directly from a highly differentiated compartment to the stem cell compartment without there being in intermediate stages (Fig 1 c). Given hierarchically structured multi-compartment cell population, we are concerned about the selection of stepwise or jumpwise de-differentiating mutant cell population in the competition with non de-differentiating resident cell population. By comparing the growth rates of different cell populations, we analyze the favorable conditions for different de-differentiation patterns to invade a tissue. We hope that our work contributes to the theoretical understanding of the emergence of de-differentiation in multicellular tissues.

## Methods

### The matrix population model for cellular hierarchy

Consider a cell population composed of *n* compartments, each of which represents a certain stage of differentiation [10, 13] (Fig. 1). For example, compartment 1 represents stem cells, and compartment *n* represents terminally differentiated cells. Each cell in compartment *i* (1 ≤ *i ≤ n −* 1) divides at rate *r_i_*. With probability *p_i_,* it divides symmetrically, giving birth to two identical cells in compartment *i* (Fig. 1 d). With probability *q_i_,* it differentiates symmetrically, generating two identical daughter cells in compartment *i* + 1. It should be pointed out that, for simplicity, asymmetric division [39, 40] (giving birth to one daughter cell in compartment *i* and the other in compartment *i* + 1) is not taken into account here, but our approaches and results are still applicable for the models with asymmetric division. The terminally differentiated cells in compartment *n* cannot divide and are removed from the tissue at rate *d*.

We use the vector 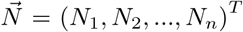 to denote the cell numbers in different compartments. Then, the hierarchically structured population dynamics composed of non de-differentiating cells can be described as a matrix population model [38]

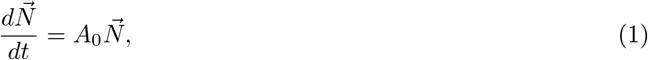

where *A*_0_ is the projection matrix which is given by

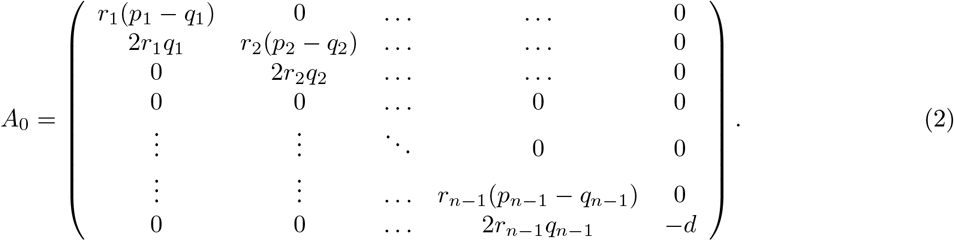

Here *r_i_*(p_*i*_ − *q*_*i*_) represents the effective self-renewal rate of compartment *i*, and 2*r*_*i*_*q*_*i*_ represents the influx rate from compartment *i* to compartment *i* + 1 due to differentiation. Let 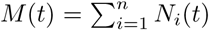 be the total cell number of the population. Note that *A*_0_ is an essentially non-negative (all the off-diagonal elements are non-negative [41]) and lower triangular matrix. According to the standard theory of matrix population models [38], the population grows exponentially in the long run, i.e.

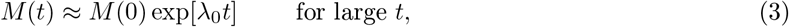

where *λ*_0_ is the real leading eigenvalue. The leading eigenvalue hence characterizes the asymptotic growth rate of the whole population, which is often used as a measure of fitness in matrix population models [42, 43]. Here, we use this fitness measure to assess whether a mutant can invade a resident population.

### Stepwise and jumpwise de-differentiation

Let us now introduce de-differentiation processes given the non de-differentiating resident cell population Eq. (1). Since it is biologically unclear how a non de-differentiating resident cell acquires the ability for de-differentiation, here we consider de-differentiation as a result of certain genetic or epigenetic alterations (jointly referred to as mutations). It is assumed that the mutant cells are provided with the additional ability of de-differentiation. More specifically, when these mutant cells divide, besides symmetric division and symmetric differentiation, they can also perform symmetric de-differentiation (Fig. 1 d) with a small probability. In principle, there are two different ways to do this: (i) stepwise de-differentiation, where cells de-differentiate to the previous compartment, and (ii) jumpwise de-differentiation, where de-differentiation happens across multiple compartments at a time. These are the most extreme cases and a mixture between them is possible.

For stepwise de-differentiation, a mutant cell in compartment *i* gives rise to two daughter cells in its adjacent upstream compartment *i* − 1 (Fig. 1 b) when de-differentiation happens. Suppose that the de-differentiation probability from compartment *i* to *i* − 1 is *δ*_*i*_. Then the influx rate from compartment *i* to *i* − 1 due to de-differentiation is given by 2*r*_*i*_*δ*_*i*_. When de-differentiation is taken into account, the self-renewal probability of each mutant cell in compartment *i* becomes 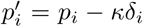, and its differentiation probability becomes 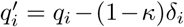, such that we have 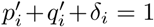. Here, the parameter *κ* (0 ≤ *κ* ≤ 1) governs how mutant cell redistributes the probabilities for self-renewal and differentiation when taking de-differentiation into account. We call *κ* the redistributing factor. De-differentiation is generally a rare event [21], we thus assume that *ρ*_*i*_ = 2*r*_*i*_*δ*_i_ ≪ 1. As the occurrence of de-differentiation for different stages of differentiation is poorly understood, for simplicity we assume that all the *ρ*_*i*_ are the same, i.e. they are independent of index *i* and denoted as *ρ* for short. In this way, the population dynamics of the stepwise de-differentiating mutant population can be modeled with a projection matrix given by

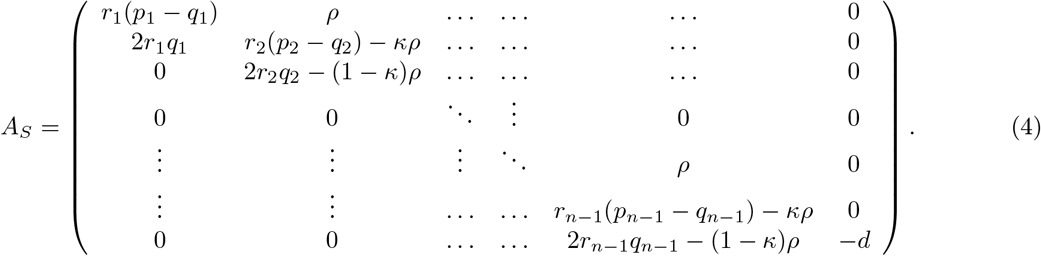

Jumpwise de-differentiation provides an alternative pattern where even highly differentiated cells can directly revert to stem cells without being in intermediate stages (Fig 1 c). Formally, it is assumed that the jumpwise de-differentiating mutant cell in compartment *n* − 1 give birth to two daughter stem cells in compartment 1 (Fig 1d). Therefore, the projection matrix is given by

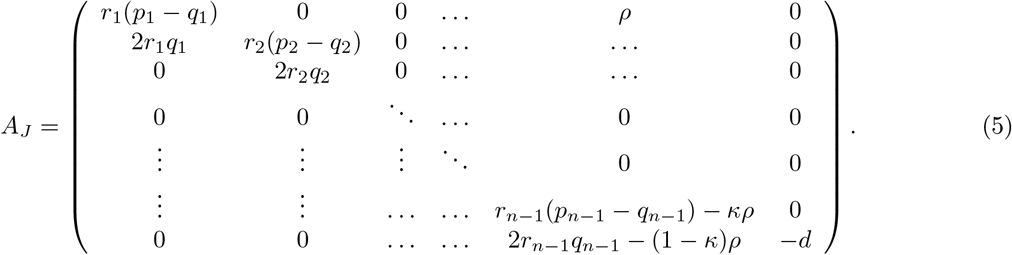

### Selection gradient for de-differentiation

In the following, we consider the competition between a non de-differentiating resident cell population and a stepwise de-differentiating mutant cell population (which is called S mutant cell population for short), as well as between a non de-differentiating resident cell population and a jumpwise de-differentiating mutant cell population (which is called *J* mutant cell population for short) by comparing their fitness measures, i.e. the leading eigenvalues λ_0_, λ_*S*_ and λ_*J*_ of *A*_0_, *A*_*S*_ and *A_J_*, respectively. Note that *ρ* is very small, such that both *A*_*S*_ and *A_J_* can be seen as matrix perturbations to *A*_0_. According to the eigenvalue perturbation method [44], we have

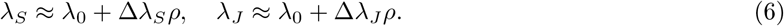

Here, Δλ_*S*_ and Δλ_*J*_ are given by

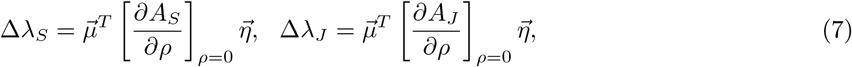

where 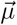 and 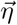 are the left and right eigenvectors associated with λ_0_ respectively (see Supplementary Information).

For a given parameter set (*r_i_*, *p_i_*, *q_i_*, *d*, *κ*), Δλ_*S*_ characterizes the selective difference between an S mutant cell population and a non de-differentiating cell population. If Δλ_*S*_ > 0, for example, the S mutant population is favored in this competition — a non de-differentiating resident cell population is invaded by an S mutant cell population. Therefore, Δλ_*S*_ corresponds to a selection gradient and acts as a comparative fitness measure of the S mutant cell population relative to the non de-differentiating resident cell population. A similar argument also applies for Δλ_*J*_. We thus term Δλ_*S*_ and Δλ_*J*_ as selection gradients of the S mutant cell population and the *J* mutant cell population, respectively. Based on these quantities, we will analyze the favorable condition for de-differentiation.

## Results

We infer whether de-differentiation leads to an increased fitness in the different cases (stepwise and jumpwise), both analytically and numerically.

Let us first focus on the null model without de-differentiation. In this case, the projection matrix *A*_0_ is a lower triangular matrix whose eigenvalues are just the diagonal elements. Suppose that the resident cell population in Eq. (1) is not shrinking, which implies that there exists at least one non-negative diagonal element in *A*_0_. In this way, the leading eigenvalue λ_0_ is the largest among all the non-negative diagonal elements of *A*_0_. Note that −*d* is always negative, such that λ_0_ is always in the form of *r_j_*_0_ (*p_j_*_0_ − *q_j_*_0_), where *j*_0_ is the compartment that maximizes this quantity.

Next, we turn to stepwise de-differentiation, cf. Eq. (4). Given λ_0_ = *r_j_*_0_ (*p_j_*_0_ − *q_j_*_0_), the selection gradient (comparative fitness) of an S mutant cell population is given by (see Supplementary Information)

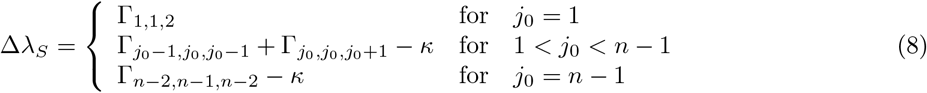

where 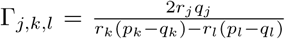. In reality, cells in different compartments do not have exactly the same effective self-renewal rate, because of internal or external noise in cellular dynamics [36]. It is thus reasonable to assume that the leading eigenvalue λ_0_ is unique, which implies that *r_j_*_0_ (*p_j_*_0_ − *q_j_*_0_) is strictly larger than any other *r_j_* (*p*_*j*_ − *q*_*j*_) for *j* ≠ *j*_0_

Thus, all the г_*j,k,l*_ in Eq. (8) are positive. In particular, for *j*_0_ = 1, Δλ_*S*_ = г_1,1,2_ is positive. In other words, an S mutant cell population is always favored in the competition with non de-differentiating resident cell population provided that stem cells have the largest effective self-renewal rate among all cell compartments. We performed exact numerical solutions to verify our theoretical approximation and find a very good agreement, see Fig. 2. Furthermore, with the increase of symmetric division probability of stem cells (*p*_1_), the selection gradient Δλ_*S*_ gradually tends to zero (Fig. 2).

**Figure 2:**
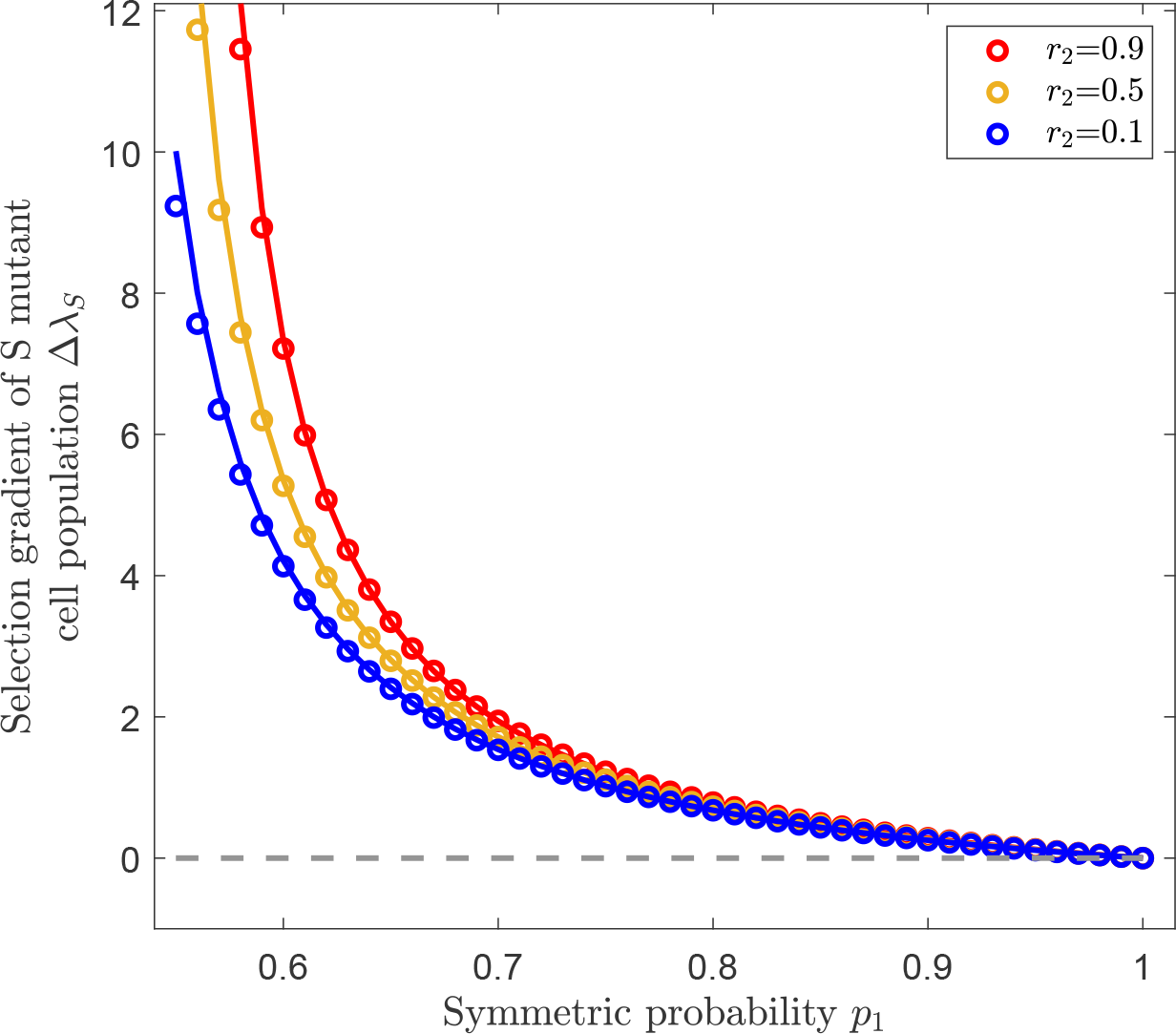
Selection for stepwise de-differentiation when the effective rate of self renewal is highest for stem cells. Illustration of the selection gradient (comparative fitness) of the S mutant cell population Δλ_*S*_ as a function of the symmetric division probability *p*_1_, provided that the stem cell compartment has the largest effective self-renewal rate, i.e. λ_0_ = *r*_1_*(p*_1_ − *q*_1_). Colored lines represent analytical approximations from Eq. (8) by using the eigenvalue perturbation method and symbols represent exact numerical solutions, which agree very well with each other. In this case, de-differentiation provides a fitness advantage for all values of *p*_1_ and *r*_2_. The parameters are *n* = 4, *κ* = 0.1, *p* = 0.001, *d =* 0.05, *p*_2_ = 0.55, *p*_3_ = 0.6, *r*_1_=0.99, *r*_3_ = 0.3. The range of *p*_1_ (0.55 < *p*_1_ < 1.0) ensures that *r*_1_(*p*_1_ − *q*_1_) is the leading eigenvalue.

Let us now consider the cases with *j*_0_ > 1. In reality, stem cells may have slower cycling time than more differentiated cells [45], or stem cell differentiation and self-renewal could be balanced (*p*_1_ = *q*_1_ = 0.5), maintaining a stable stem cell population [10]. Thus, sometimes it is more likely for other cell compartments (*j*_0_ > 1) to have the largest effective self-renewal rate. From Eq. (8) we can see that Δλ_*S*_ is a linear combination of г_*j,k,l*_ and *κ*. It is interesting to see that Δλ_*S*_ is negatively correlated with *κ*. Note that *κ* is the redistributing factor that characterizes how the introduction of de-differentiation reshapes the probabilities for self-renewal and differentiation. For *κ* = 0, Δλ_*S*_ is positive. With an increase of *κ*, Δλ_*S*_ could become negative. Fig. 3 typically shows two scenarios of Δλ_*S*_: either it is always larger than zero for any *κ*, or it changes from positive to negative at some critical point 0 < *κ*\* < 1. For the latter scenario, Fig 4 illustrates how Δλ_*S*_ changes with both the redistributing factor *κ* and the symmetric division probability *p*_2_ provided that compartment 2 has the largest effective self-renewal rate (*j*_0_ =2). It is shown that with the increase of *p*_2_, the critical value *κ*\* decreases, which means it is getting less likely for the S mutant cell population to be favored. Note that г_*j,k,l*_ represents the effect of cellular hierarchy on de-differentiation, and *κ* represents how de-differentiation reshapes the cellular division patterns. Therefore, the selection of de-differentiation is a combined result of cellular hierarchy and de-differentiation pattern.

**Figure 3:**
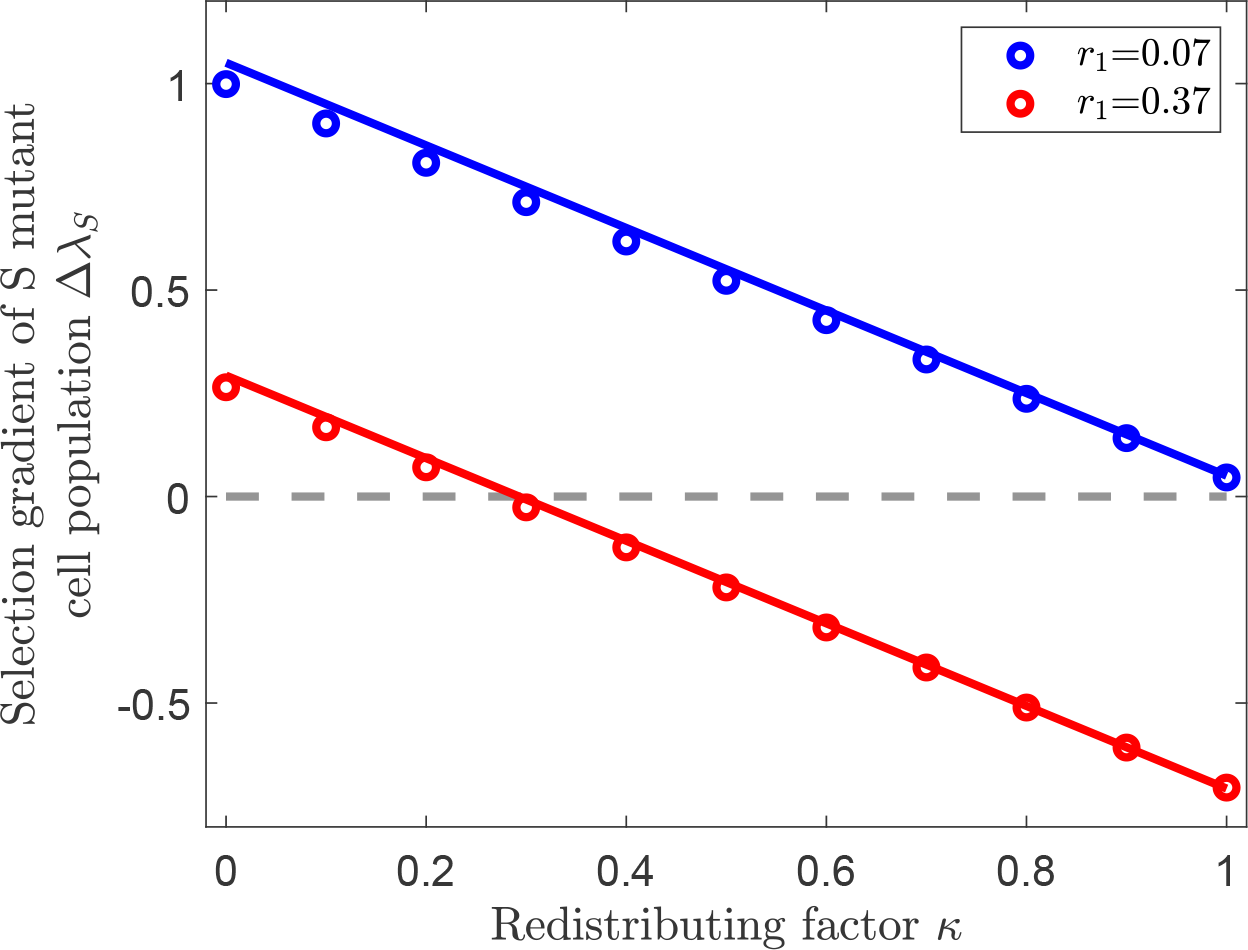
Selection for stepwise de-differentiation when the effective rate of self renewal is highest in compartment 2. Illustration of the selection gradient (comparative fitness) of the S mutant cell population Δλ_*S*_ as a function of redistributing factor *κ* provided that λ_0_ = *r*_2_(*p*_2_ − *q*_2_). Colored lines represent eigenvalue perturbation results from Eq. (8) and symbols represent exact numerical solutions. There are two different scenarios: For 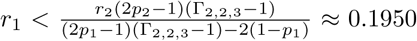, Δλ_*S*_ is always positive (blue color); For *r*_1_ > 0.1950, Δλ_*S*_ changes from positive to negative with the increase of *κ* (red color). The background parameters are *n* = 4, *ρ* = 0.01, *d =* 0.05, *p*_1_ = 0.5, *p*_2_ = 0.95, *p*_3_ = 0.55, *r*_2_=0.44, *r*_3_=0.17.

**Figure 4:**
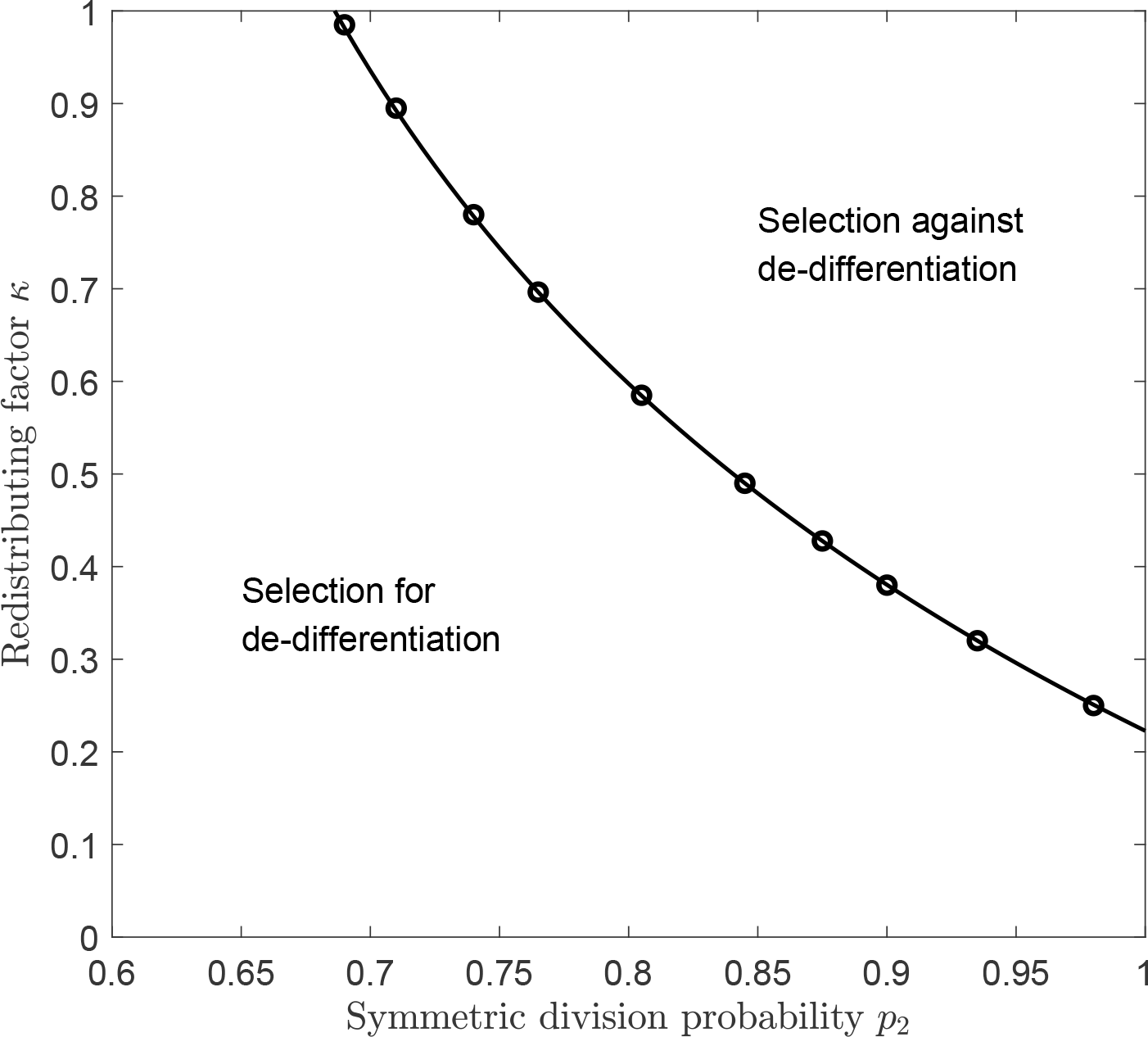
Selection for stepwise de-differentiation in a landscape composed of the symmetric division probability *p*_2_ and redistributing factor *κ* when the effective rate of self renewal is highest in compartment 2. The curve represents the boundary with Δλ_*S*_ = 0, which is generated by the eigenvalue perturbation approximation in Eq. (8). The symbols represent the exact numerical solutions for Δλ_*S*_ = 0. The parameters are *n* = 4, *ρ* = 0.01, *r*_1_ = 0.0885, *r*_2_ = 0.4145, *r*_3_ = 0.5555, *p*_1_ = 0.4723, *p*_3_ = 0.0727, *d* = 0.005.

We now turn our attention to the selection gradient (comparative fitness) of the *J* mutant cell population, which is given by (see Supplementary Information for mathematical details)

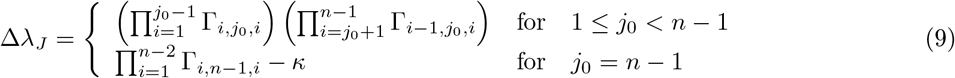

Similar to Eq. (8), here all the г_*j,k,l*_ in Eq. (9) are positive. For the cases with 1 ≤ *j*_0_ < *n* − 1, in particular, Δλ_*J*_ is always positive (Fig. 5), i.e. the *J* mutant cell population is favored in all these cases. On the other hand, for *j*_0_ = *n* − 1, Δλ_*J*_ is negatively correlated with the redistributing factor *κ* which is similar to Δλ_*S*_ (Fig. 6).

**Figure 5:**
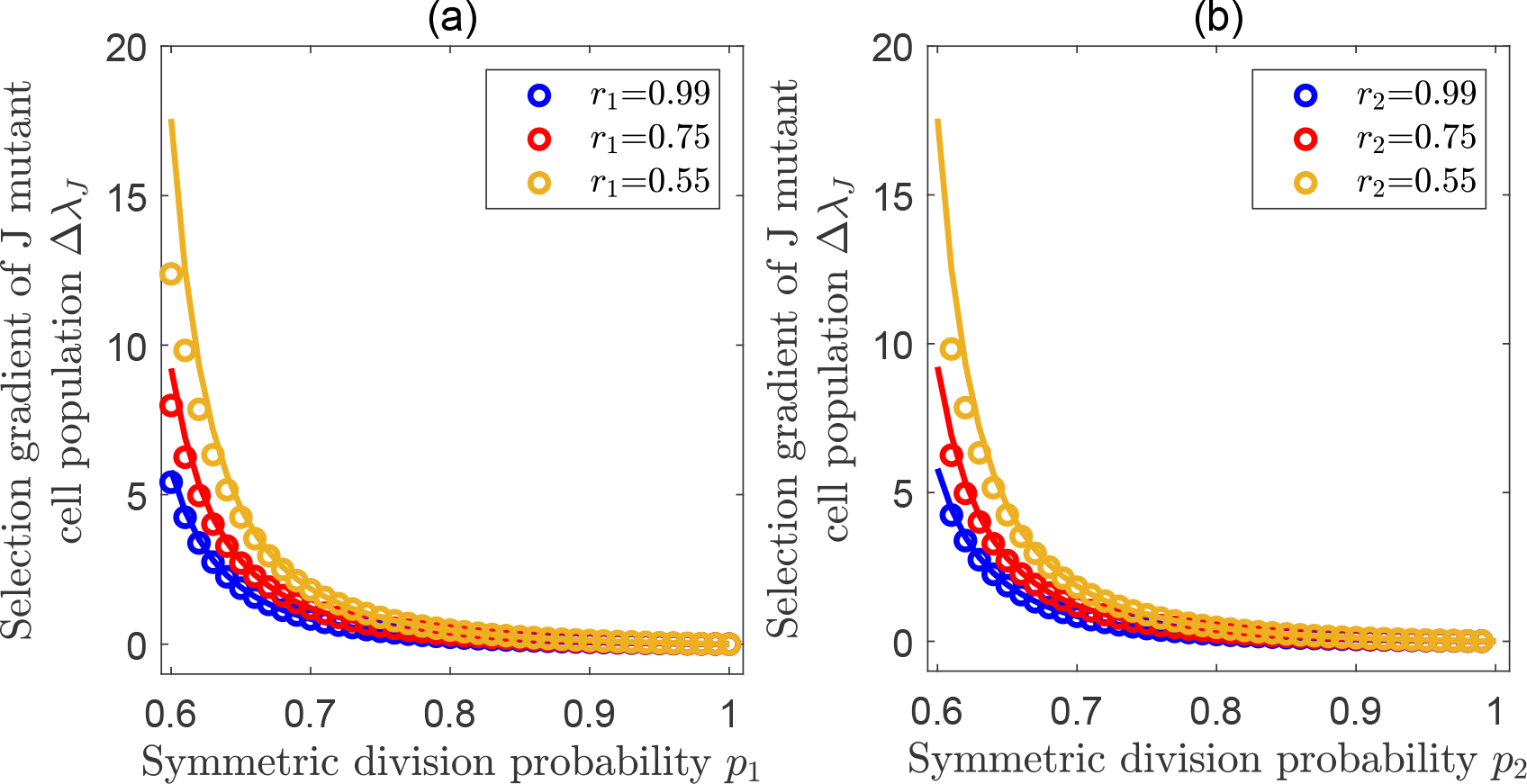
Selection for jumpwise de-differentiation. Illustrations of the selection gradient (comparative fitness) of the *J* mutant cell population Δλ_*J*_ for the cases *j*_0_ = 1 (a) and *j*_0_ = 2 (b). In both panels, colored lines represent analytical approximations from Eq. (9) by using eigenvalue perturbation and symbols represent exact numerical solutions. **(a)** Δλ_*J*_ as a function of *p*_1_ provided that compartment 1 has the largest effective self renewal rate, λ_0_ = *r*_1_(*p*_1_ − *q*_1_) for *p*_2_ = 0.55. **(b)** Δλ_*J*_ as a function of *p*_2_ provided compartment 2 has the largest effective self renewal rate, λ_0_ = *r*_2_*(p*_2_ − *q*_2_) for *p*_1_ = 0.55 (joint parameters *n* = 4, *κ* = 0.1, *ρ* = 0.01, *d* = 0.05, *p*_3_ = 0.6, *r*_1_= 0.2, *r*_3_ = 0.3).

**Figure 6:**
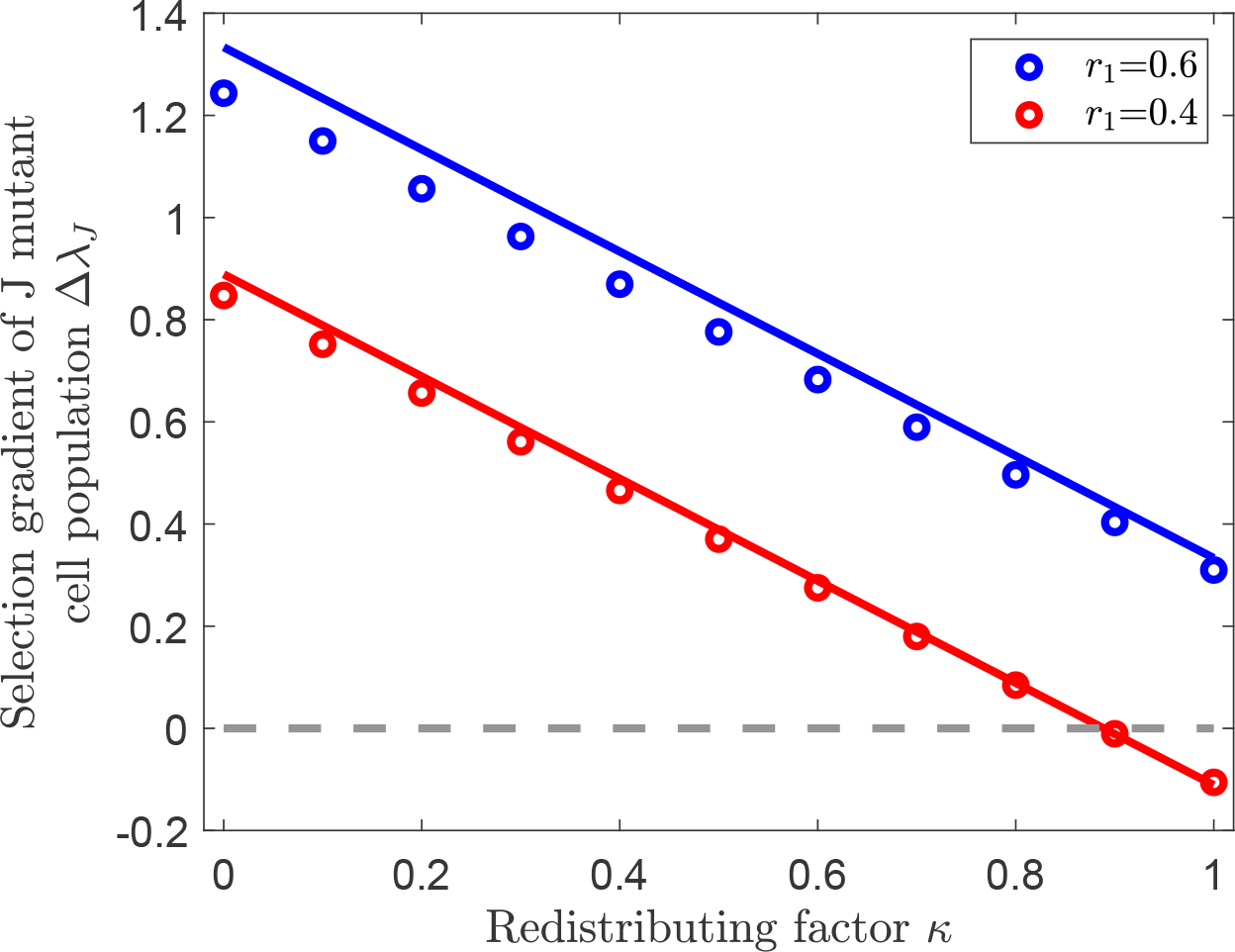
Selection for jumpwise de-differentiation when the effective rate of self renewal is highest in compartment 3. Illustration of the selection gradient Δλ_*J*_ as a function of the redistributing factor *κ* provided that λ_0_ = *r*_3_(*p*_3_ − *q*_3_). Colored lines represent eigenvalue perturbation results in Eq. (9) and symbols represent exact numerical solutions. There are two different scenarios: For 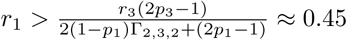, Δλ_*J*_ is always positive (blue color). For *r*_1_ < 0.45, Δλ_*S*_ is changed from positive to negative with the increase of *κ* (red color). The background parameters are *n* = 4, *ρ* = 0.01, *d =* 0.05, *p*_1_ = 0.5, *p*_2_ = 0.65, *p*_3_ = 0.85, *r*_2_ = 0.4, *r*_3_=0.6.

A comparison between Eqs. (8) and (9) reveals some important differences between stepwise and jumpwise de-differentiation patterns. First of all, jumpwise de-differentiation provides a much wider range of favorable condition for de-differentiation than stepwise de-differentiation in the sense that Δλ_*J*_ is always positive for any 1 ≤ *j*_0_ < *n* − 1, but Δλ_*S*_ is always positive only for *j*_0_ = 1. Secondly, Δλ_*S*_ only depends on the parameters related to the neighborhood compartments of *j*_0_, but Δλ_*J*_ depends on the parameters related to all compartments, ranging from the stem cell stage to the stage where de-differentiation occurs. This implies that, the total number of compartment does matter in the jumpwise case, but not in the stepwise case. In other words, stepwise de-differentiation utilizes the local structure around the compartment with the largest effective self-renewal rate, whereas jumpwise de-differentiation utilizes the global structure throughout the multi-compartment hierarchy.

## Discussion

In this study, we have explored the adaptive significance of de-differentiation in hierarchical multi-compartment structured cell populations. By establishing a matrix population model, we study the competition between resident hierarchical structured cell populations without de-differentiation and mutant cell populations with different modes of de-differentiation.

In principle, there are two main factors that could influence the selection of de-differentiation: cellular hierarchy and the de-differentiation pattern. Cellular hierarchy refers e.g. to the number of cell compartments, the inherent cell division pattern, or the cell division rate. These correspond to the parameter landscape of (*n*, *p*_i_, *q*_i_, *r*_i_) in our model. The de-differentiation pattern refers to different modes of de-differentiation (stepwise or jumpwise), as well as how de-differentiation reshapes the division pattern in the cellular hierarchy (corresponding to *κ* in our model). Interestingly, our results show that the selection gradients for de-differentiation (Δλ_*S*_ and Δλ_*J*_) can generally be decomposed into a sum of a cellular hierarchy part and a de-differentiation part, showing that the selection of de-differentiation is a result of the linear combinations of these two factors.

Among all factors in the cellular hierarchy, the most important one is which of the cell compartments has the largest effective self-renewal rate. This is because de-differentiation is more likely to be favored when earlier compartments have the largest effective self-renewal rate. For example, in the stepwise case, de-differentiation is favored provided that the stem cell has the largest effective self-renewal rate. In the jumpwise case, de-differentiation is favored in all cases except when the latest divisible cell compartment has the largest effective self-renewal rate.

Given all the factors in the cellular hierarchy, we are most concerned about how different de-differentiation patterns shape the evolution of de-differentiation. In particular the redistributing factor, i.e. the effect of dedifferentiation on self-renewal and differentiation probabilities greatly influences the selection condition. Our results suggest that de-differentiation is more likely to be favored if there is less effect on self-renewal than on differentiation. In addition, the de-differentiation mode (stepwise or jumpwise) has enormous implications for the selection condition. Our results suggest that de-differentiation is more likely to be favored in the jumpwise case than in the stepwise case. However, jumpwise de-differentiation seems to be biologically much more difficult to achieve, the overall incidence of it would still be very low. Perhaps an example of the differences between stepwise and jumpwise de-differentiation and the implications of the subsequent disease behavior can be illustrated by various types of leukemia. As already mentioned, MLL-AF9 expression in a committed progenitor cells can lead to the development of leukemic stem cells that can result in disease transmission across mice [27, 46]. In general MLL expression is associated with a poor prognosis in acute myeloid leukemia [47, 48]. This may be an example of jumpwise de-differentiation. In contrast, acute promyelocytic leukemia (APL) is an example of acute leukemia that is highly curable [49]. It is therefore possible that in this disease, stepwise de-differentiation - or a situation where a mutant cell can stick in a compartment without differentiating, similar to a stem cell - is occurring that in part makes the disease still potentially curable.

Note that the presented study is based on a matrix population model with constant elements, which in principle captures an exponential growing population. Even though there are still uncertainties regarding the growth patterns of cell populations in different contexts (cancer or normal, solid or hematologic tumor, in vivo or in vitro) [7, 50] and exponential growth is often considered to be unable to capture the biological processes in reality, exponential-like growth models are widely used as default models to describe growing cell populations, especially in early cancer development [21, 51-53]. We followed this idea and used it as a starting point to explore the adaptive significance of de-differentiation. In the future, more complex biological mechanisms such as density-dependent population growth could be taken into account in models for de-differentiation. Moreover, while the hierarchical architecture of tissues is considered to have been selected to minimize the risk of retention of mutations, the risk of acquisition of stem cell line properties by the large population of progenitor cells introduces new dynamics - perhaps in such a scenario two additional considerations could reduce the risk of cancer - namely the low probability that specific mutations lead to acquisition of stem cell like behavior or the average survival of progenitor cells may be low enough to prevent the acquisition of the additional mutations needed to reach the full cancer phenotype. This could be an extension of this work in future.

## References

[1] Michor F, Nowak MA, Frank SA, Iwasa Y. Stochastic elimination of cancer cells. Proceedings of the Royal Society of London B: Biological Sciences. 2003;270(1528):2017–2024.

[2] Nowak MA, Michor F, Iwasa Y. The linear process of somatic evolution. Proceedings of the National Academy of Sciences. 2003;100(25):14966–14969.

[3] Dick JE. Stem cell concepts renew cancer research. Blood. 2008;112(13):4793–4807.

[4] Fichelson P, Audibert A, Simon F, Gho M. Cell cycle and cell-fate determination in Drosophila neural cell lineages. Trends in Genetics. 2005;21(7):413–420.

[5] Michor F, Hughes TP, Iwasa Y, Branford S, Shah NP, Sawyers CL, et al. Dynamics of chronic myeloid leukaemia. Nature. 2005;435(7046):1267.

[6] Dzierzak E, Speck NA. Of lineage and legacy: the development of mammalian hematopoietic stem cells. Nature Immunology. 2008;9(2):129.

[7] Johnston MD, Edwards CM, Bodmer WF, Maini PK, Chapman SJ. Mathematical modeling of cell population dynamics in the colonic crypt and in colorectal cancer. Proceedings of the National Academy of Sciences. 2007;104(10):4008–4013.

[8] Dingli D, Traulsen A, Pacheco JM. Compartmental architecture and dynamics of hematopoiesis. PLoS ONE. 2007;2(4):e345.

[9] Takizawa H, Regoes RR, Boddupalli CS, Bonhoeffer S, Manz MG. Dynamic variation in cycling of hematopoietic stem cells in steady state and inflammation. Journal of Experimental Medicine. 2011;208(2):273–284.

[10] Werner B, Dingli D, Lenaerts T, Pacheco JM, Traulsen A. Dynamics of mutant cells in hierarchical organized tissues. PLoS Computational Biology. 2011;7(12):e1002290.

[11] Rodriguez-Brenes IA, Wodarz D, Komarova NL. Minimizing the risk of cancer: tissue architecture and cellular replication limits. Journal of The Royal Society Interface. 2013;10(86):20130410.

[12] Alvarado C, Fider NA, Wearing HJ, Komarova NL. Optimizing homeostatic cell renewal in hierarchical tissues. PLoS Computational Biology. 2018;14(2):e1005967.

[13] Böttcher MA, Dingli D, Werner B, Traulsen A. Replicative cellular age distributions in compartmentalized tissues. Journal of The Royal Society Interface. 2018;15(145):20180272.

[14] Reya T, Morrison SJ, Clarke MF, Weissman IL. Stem cells, cancer, and cancer stem cells. Nature. 2001;414(6859):105.

[15] Jordan CT, Guzman ML, Noble M. Cancer stem cells. New England Journal of Medicine. 2006;355(12):1253–1261.

[16] Altrock PM, Liu LL, Michor F. The mathematics of cancer: integrating quantitative models. Nature Reviews Cancer. 2015;15(12):730.

[17] Tata PR, Mou H, Pardo-Saganta A, Zhao R, Prabhu M, Law BM, et al. Dedifferentiation of committed epithelial cells into stem cells in vivo. Nature. 2013;503(7475):218.

[18] Easwaran H, Tsai HC, Baylin SB. Cancer epigenetics: tumor heterogeneity, plasticity of stem-like states, and drug resistance. Molecular Cell. 2014;54(5):716–727.

[19] Tetteh PW, Farin HF, Clevers H. Plasticity within stem cell hierarchies in mammalian epithelia. Trends in Cell Biology. 2015;25(2):100–108.

[20] Chaffer CL, Brueckmann I, Scheel C, Kaestli AJ, Wiggins PA, Rodrigues LO, et al. Normal and neoplastic nonstem cells can spontaneously convert to a stem-like state. Proceedings of the National Academy of Sciences. 2011;108(19):7950–7955.

[21] Gupta PB, Fillmore CM, Jiang G, Shapira SD, Tao K, Kuperwasser C, et al. Stochastic state transitions give rise to phenotypic equilibrium in populations of cancer cells. Cell. 2011;146(4):633–644.

[22] Meacham CE, Morrison SJ. Tumour heterogeneity and cancer cell plasticity. Nature. 2013;501(7467):328.

[23] Yang G, Quan Y, Wang W, Fu Q, Wu J, Mei T, et al. Dynamic equilibrium between cancer stem cells and non-stem cancer cells in human SW620 and MCF-7 cancer cell populations. British Journal of Cancer. 2012;106(9):1512.

[24] Quintana E, Shackleton M, Foster HR, Fullen DR, Sabel MS, Johnson TM, et al. Phenotypic heterogeneity among tumorigenic melanoma cells from patients that is reversible and not hierarchically organized. Cancer cell. 2010;18(5):510–523.

[25] Dorantes-Acosta E, Pelayo R. Lineage switching in acute leukemias: a consequence of stem cell plasticity? Bone Marrow Research. 2012;2012.

[26] Passegué E, Weisman IL. Leukemic stem cells: where do they come from? Stem Cell Reviews. 2005;1(3):181–188.

[27] Krivtsov AV, Twomey D, Feng Z, Stubbs MC, Wang Y, Faber J, et al. Transformation from committed progenitor to leukaemia stem cell initiated by MLL–AF9. Nature. 2006;442(7104):818.

[28] Haeno H, Levine RL, Gilliland DG, Michor F. A progenitor cell origin of myeloid malignancies. Proceedings of the National Academy of Sciences. 2009;106(39):16616–16621.

[29] Leder K, Pitter K, LaPlant Q, Hambardzumyan D, Ross BD, Chan TA, et al. Mathematical modeling of PDGF-driven glioblastoma reveals optimized radiation dosing schedules. Cell. 2014;156(3):603–616.

[30] Jilkine A, Gutenkunst RN. Effect of dedifferentiation on time to mutation acquisition in stem cell-driven cancers. PLoS Computational Biology. 2014;10(3):e1003481.

[31] Mahdipour-Shirayeh A, Kaveh K, Kohandel M, Sivaloganathan S. Phenotypic heterogeneity in modeling cancer evolution. PLoS ONE. 2017;12(10):e0187000.

[32] Wodarz D. Effect of cellular de-differentiation on the dynamics and evolution of tissue and tumor cells in mathematical models with feedback regulation. Journal of Theoretical Biology. 2018;448:86–93.

[33] dos Santos RV, da Silva LM. A possible explanation for the variable frequencies of cancer stem cells in tumors. PLoS ONE. 2013;8(8):e69131.

[34] Niu Y, Wang Y, Zhou D. The phenotypic equilibrium of cancer cells: From average-level stability to path-wise convergence. Journal of Theoretical Biology. 2015;386:7–17.

[35] Zhou JX, Pisco AO, Qian H, Huang S. Nonequilibrium population dynamics of phenotype conversion of cancer cells. PLoS ONE. 2014;9(12):e110714.

[36] Zhou D, Wang Y, Wu B. A multi-phenotypic cancer model with cell plasticity. Journal of Theoretical Biology. 2014;357:35–45.

[37] Chen X, Wang Y, Feng T, Yi M, Zhang X, Zhou D. The overshoot and phenotypic equilibrium in characterizing cancer dynamics of reversible phenotypic plasticity. Journal of Theoretical Biology. 2016;390:40–49.

[38] Caswell H. Matrix Population Models. John Wiley & Sons, Ltd; 2006.

[39] Dingli D, Traulsen A, Michor F. (A) symmetric stem cell replication and cancer. PLoS Computational Biology. 2007;3(3):e53.

[40] Hu Z, Fu YX, Greenberg AJ, Wu CI, Zhai W. Age-dependent transition from cell-level to population-level control in murine intestinal homeostasis revealed by coalescence analysis. PLoS Genetics. 2013;9(2):e1003326.

[41] Cohen JE. Convexity of the dominant eigenvalue of an essentially nonnegative matrix. Proceedings of the American Mathematical Society. 1981;81(4):657–658.

[42] Metz JA, Nisbet RM, Geritz SA. How should we define ‘fitness’ for general ecological scenarios? Trends in Ecology & Evolution. 1992;7(6):198–202.

[43] Pichugin Y, Peña J, Rainey PB, Traulsen A. Fragmentation modes and the evolution of life cycles. PLoS Computational Biology. 2017;13(11):e1005860.

[44] Trefethen LN, Bau III D. Numerical Linear Algebra. vol. 50. Philadelphia, PA: SIAM; 1997.

[45] Fillmore CM, Kuperwasser C. Human breast cancer cell lines contain stem-like cells that self-renew, give rise to phenotypically diverse progeny and survive chemotherapy. Breast Cancer Research. 2008;10(2):R25.

[46] Dong F, Bai H, Wang X, Zhang S, Wang Z, Xie M, et al. Mouse acute leukemia develops independent of self-renewal and differentiation potentials in hematopoietic stem and progenitor cells. Blood Advances. 2019;3(3):419–431.

[47] Stavropoulou V, Kaspar S, Brault L, Sanders MA, Juge S, Morettini S, et al. MLL-AF9 expression in hematopoietic stem cells drives a highly invasive AML expressing EMT-related genes linked to poor outcome. Cancer Cell. 2016;30(1):43–58.

[48] Scholl C, Schlenk RF, Eiwen K, Dohner H, Frohling S, Dohner K, et al. The prognostic value of MLL-AF9 detection in patients with t (9; 11)(p22; q23)-positive acute myeloid leukemia. Haematologica. 2005;90(12):1626–1634.

[49] Werner B, Gallagher RE, Paietta EM, Litzow MR, Tallman MS, Wiernik PH, et al. Dynamics of leukemia stem-like cell extinction in acute promyelocytic leukemia. Cancer Research. 2014;74(19):5386–5396.

[50] Gerlee P. The model muddle: in search of tumour growth laws. Cancer Research. 2013; p. canres–4355.

[51] Rodriguez-Brenes IA, Komarova NL, Wodarz D. Tumor growth dynamics: insights into evolutionary processes. Trends in Ecology & Evolution. 2013;28(10):597–604.

[52] Williams MJ, Werner B, Barnes CP, Graham TA, Sottoriva A. Identification of neutral tumor evolution across cancer types. Nature Genetics. 2016;48(3):238.

[53] Talkington A, Durrett R. Estimating tumor growth rates in vivo. Bulletin of Mathematical Biology. 2015;77(10):1934–1954.

