## Supplementary material for "The invasion of de-differentiating cancer cells into hierarchical tissues": Supplimentary Information

#### Calculation of selection gradients $\Delta\lambda_S$ and $\Delta\lambda_J$

Consider a matrix population model [1]

$$\frac{d\vec{N}}{dt} = A_0\vec{N}, \tag{1}$$

where the projection matrix is given by

$$A_0 = \begin{pmatrix} a_{11} & 0 & 0 & \dots & \dots & 0 \\ a_{21} & a_{22} & 0 & \dots & \dots & 0 \\ 0 & a_{32} & 0 & \dots & \dots & 0 \\ 0 & 0 & \ddots & \dots & 0 & 0 \\ \vdots & \vdots & \vdots & \ddots & 0 & 0 \\ \vdots & \vdots & \dots & \dots & a_{n-1.n-1} & 0 \\ 0 & 0 & \dots & \dots & a_{n,n-1} & a_{nn} \end{pmatrix}. \tag{2}$$

Note that $A_0$ is actually an abstraction of the $A_0$ in the main text where

$$a_{ii} = \begin{cases} r_i(p_i - q_i) & \text{for} \quad 1 \le i < n-1 \\ -d & \text{for} \quad i = n \end{cases} \tag{3}$$

$$a_{i+1,i} = 2r_iq_i. \tag{4}$$

$A_0$ is a a lower triangular matrix, whose eigenvalues are just the diagonal elements. Suppose the leading eigenvalue of $A_0$ is $\lambda_0 = a_{j_0j_0}$, then we can calculate the associated left eigenvector as

$$\vec{\mu} = \frac{1}{\prod_{i=1}^{j_0-1}\left(\frac{a_{j_0j_0}-a_{ii}}{a_{i+1,i}}\right)}\left(1, \frac{a_{j_0j_0}-a_{11}}{a_{21}}, \frac{(a_{j_0j_0}-a_{11})(a_{j_0j_0}-a_{22})}{a_{21}a_{32}}, ..., \prod_{i=1}^{j_0-1}\left(\frac{a_{j_0j_0}-a_{ii}}{a_{i+1,i}}\right), 0, ..., 0\right)^T \tag{5}$$

and its right eigenvalue as (here $\vec{\mu}^T\vec{\eta} = 1$)

$$\vec{\eta} = \left(0, ..., 0, 1, \frac{a_{j_0+1,j_0}}{a_{j_0,j_0} - a_{j_0+1,j_0+1}}, ..., \prod_{i=j_0}^{n-1}\frac{a_{i+1,i}}{a_{j_0,j_0} - a_{i+1,i+1}}\right)^T. \tag{6}$$

Suppose that the population in Eq. (1) is not shrinking, which implies that there exists at least one non-negative diagonal element in $A_0$. In this way, the leading eigenvalue $\lambda_0$ is the largest among all the non-negative diagonal elements of $A_0$. Notice that $a_{nn} = -d$ is always negative, it cannot be the leading eigenvalue, the possible value of $j_0$ hence can only range from 1 to $n-1$.

Consider two different types of matrix perturbation to $A_0$ as follows ($\rho \ll 1$):

$$A_S = \begin{pmatrix} a_{11} & \rho & \dots & \dots & \dots & 0 \\ a_{21} & a_{22} - \kappa\rho & \dots & \dots & \dots & 0 \\ 0 & a_{32} - (1-\kappa)\rho & \dots & \dots & \dots & 0 \\ 0 & 0 & \ddots & \dots & 0 & 0 \\ \vdots & \vdots & \vdots & \ddots & \rho & 0 \\ \vdots & \vdots & \dots & \dots & a_{n-1,n-1} - \kappa\rho & 0 \\ 0 & 0 & \dots & \dots & a_{n,n-1} - (1-\kappa)\rho & a_{nn} \end{pmatrix} \tag{7}$$

and

$$A_J = \begin{pmatrix} a_{11} & 0 & 0 & \dots & \rho & 0 \\ a_{21} & a_{22} & 0 & \dots & \dots & 0 \\ 0 & a_{32} & 0 & \dots & \dots & 0 \\ 0 & 0 & \ddots & \dots & 0 & 0 \\ \vdots & \vdots & \vdots & \ddots & 0 & 0 \\ \vdots & \vdots & \dots & \dots & a_{n-1.n-1} - \kappa\rho & 0 \\ 0 & 0 & \dots & \dots & a_{n,n-1} - (1-\kappa)\rho & a_{nn} \end{pmatrix} \tag{8}$$

correspond to the stepwise and jumpwise de-differentiation cases, respectively. $\lambda_S$ and $\lambda_J$ are the leading eigenvalues of $A_S$ and $A_J$ respectively.

According to perturbation theory for eigenvalue problems [2], we have

$$\lambda_S \approx \lambda_0 + \Delta\lambda_S \rho, \quad \lambda_J \approx \lambda_0 + \Delta\lambda_J \rho. \tag{9}$$

Here, $\Delta\lambda_S$ and $\Delta\lambda_J$ are given by

$$\Delta\lambda_S = \vec{\mu}^T \left[\frac{\partial A_S}{\partial \rho}\right]_{\rho=0} \vec{\eta}, \;\; \Delta\lambda_J = \vec{\mu}^T \left[\frac{\partial A_J}{\partial \rho}\right]_{\rho=0} \vec{\eta}, \tag{10}$$

where $\vec{\mu}$ and $\vec{\eta}$ are the left and right eigenvectors associated with $\lambda_0$ respectively. Note that

$$\left[\frac{\partial A_S}{\partial \epsilon}\right]_{\epsilon=0} = \begin{pmatrix} 0 & 1 & \dots & \dots & \dots & 0 \\ 0 & -\kappa & \dots & \dots & \dots & 0 \\ 0 & -(1-\kappa) & \dots & \dots & \dots & 0 \\ 0 & 0 & \ddots & \dots & 0 & 0 \\ \vdots & \vdots & \vdots & \ddots & \rho & 0 \\ \vdots & \vdots & \dots & \dots & -\kappa & 0 \\ 0 & 0 & \dots & \dots & -(1-\kappa) & 0 \end{pmatrix} \tag{11}$$

and

$$\left[\frac{\partial A_J}{\partial \epsilon}\right]_{\epsilon=0} = \begin{pmatrix} 0 & 0 & 0 & \dots & 1 & 0 \\ 0 & 0 & 0 & \dots & \dots & 0 \\ 0 & 0 & 0 & \dots & \dots & 0 \\ 0 & 0 & \ddots & \dots & 0 & 0 \\ \vdots & \vdots & \vdots & \ddots & 0 & 0 \\ \vdots & \vdots & \dots & \dots & -\kappa & 0 \\ 0 & 0 & \dots & \dots & -(1-\kappa) & 0 \end{pmatrix}. \tag{12}$$

Then we have

$$\Delta\lambda_S = \begin{cases} \frac{a_{21}}{a_{11} - a_{22}} & \text{for} \quad j_0 = 1 \\ \frac{a_{j_0,j_0-1}}{a_{j_0,j_0} - a_{j_0-1,j_0-1}} + \frac{a_{j_0+1,j_0}}{a_{j_0,j_0} - a_{j_0+1,j_0+1}} - \kappa & \text{for} \quad 1 < j_0 < n-1 \\ \frac{a_{n-1,n-2}}{a_{n-1,n-1} - a_{n-2,n-2}} - \kappa & \text{for} \quad j_0 = n-1 \end{cases} \tag{13}$$

and

$$\Delta\lambda_J = \begin{cases} \frac{\prod_{i=1}^{n-2} a_{i+1,i}}{\prod_{i=1,i\neq j_0}^{n-1}(a_{j_0,j_0}-a_{ii})} & \text{for} \quad 1 \leq j_0 < n-1 \\ \frac{\prod_{i=1}^{n-2} a_{i+1,i}}{\prod_{i=1,i\neq j_0}^{n-1}(a_{j_0,j_0}-a_{ii})} - \kappa & \text{for} \quad j_0 = n-1 \end{cases}. \tag{14}$$

In particular, by substituting Eqs. (3) and (4) into the above results, we obtain

$$\Delta\lambda_S = \begin{cases} \frac{2r_1q_1}{r_1(p_1-q_1)-r_2(p_2-q_2)} & \text{for} \quad j_0 = 1 \\ \frac{2r_{j_0-1}q_{j_0-1}}{r_{j_0}(p_{j_0}-q_{j_0})-r_{j_0-1}(p_{j_0-1}-q_{j_0-1})} + \frac{2r_{j_0}q_{j_0}}{r_{j_0}(p_{j_0}-q_{j_0})-r_{j_0+1}(p_{j_0+1}-q_{j_0+1})} - \kappa & \text{for} \quad 1 < j_0 < n-1 \\ \frac{2r_{n-2}q_{n-2}}{r_{n-1}(p_{n-1}-q_{n-1})-r_{n-2}(p_{n-2}-q_{n-2})} - \kappa & \text{for} \quad j_0 = n-1 \end{cases} \tag{15}$$

and

$$\Delta\lambda_J = \begin{cases} \frac{\prod_{i=1}^{n-2}(2r_iq_i)}{\prod_{i=1,i\neq j_0}^{n-1}(r_{j_0}(p_{j_0}-q_{j_0})-r_i(p_i-q_i))}. & \text{for} \quad 1 \leq j_0 < n-1 \\ \frac{\prod_{i=1}^{n-2}(2r_iq_i)}{\prod_{i=1,i\neq j_0}^{n-1}(r_{j_0}(p_{j_0}-q_{j_0})-r_i(p_i-q_i))} - \kappa. & \text{for} \quad j_0 = n-1 \end{cases}. \tag{16}$$

If we defne $\Gamma_{j,k,l} = \frac{2r_jq_j}{r_k(p_k-q_k)-r_l(p_l-q_l)}$, then the above results can be simplified to

$$\Delta\lambda_S = \begin{cases} \Gamma_{1,1,2} & \text{for} \quad j_0 = 1 \\ \Gamma_{j_0-1,j_0,j_0-1} + \Gamma_{j_0,j_0,j_0+1} - \kappa & \text{for} \quad 1 < j_0 < n-1 \\ \Gamma_{n-2,n-1,n-2} - \kappa & \text{for} \quad j_0 = n-1 \end{cases} \tag{17}$$

and

$$\Delta\lambda_J = \begin{cases} \left(\prod_{i=1}^{j_0-1}\Gamma_{i,j_0,i}\right)\left(\prod_{i=j_0+1}^{n-1}\Gamma_{i-1,j_0,i}\right) & \text{for} \quad 1 \leq j_0 < n-1 \\ \prod_{i=1}^{n-2}\Gamma_{i,n-1,i} - \kappa & \text{for} \quad j_0 = n-1 \end{cases}, \tag{18}$$

which completes the calculation of the selection gradients $\Delta\lambda_S$ and $\Delta\lambda_J$ in the main text.

### References


[1] Hal Caswell. *Matrix Population Models*. John Wiley & Sons, Ltd, 2006.

[2] Lloyd N Trefethen and David Bau III. *Numerical Linear Algebra*, volume 50. Philadelphia, PA: SIAM, 1997.
